# Discovery of Chirally-dependent Protein O-2-Hydroxyglutarylation by D2HG and L2HG

**DOI:** 10.1101/2025.01.24.634716

**Authors:** Zheng Zhang, Yi-Kai Liu, Zhuojun Luo, Meng-Ju Wu, Claudia N. Evans, Zihan Qu, Fanglei Xue, Zhong-Yin Zhang, Elizabeth I. Parkinson, Nabeel Bardeesy, W. Andy Tao

## Abstract

Mutations in isocitrate dehydrogenase 1 (IDH1) and IDH2 are common in multiple types of human cancer, leading to the accumulation of D-2-hydroxyglutarate (D2HG) and the promotion of tumorigenesis^1^. Here we discovered a novel *O*-2- hydroxyglutarylation by D2HG using chemical proteomics and further revealed distinct chiral preferences for D/L2HG modifications. Notably, we identified two kinases, MRCKA and SLK, modified by D2HG and L2HG respectively, and detected reduced phosphorylation of their substrates, suggesting an inhibitory effect of D/L 2HG modifications on the kinases’ activity.

---

Recent evidence indicates that oncometabolite D2HG can be produced through alternative enzymatic pathways in cells aside from IDH mutations^2^. This phenomenon raises questions about its broader biological roles. We suspected that D2HG may also function through post-translational modifications (PTMs) of target proteins, potentially presenting novel PTMs with therapeutic relevance. To investigate potential covalent protein modifications by D2HG, we first conducted LC-MS screening on mass shifts of tryptic peptides using a wild-type human intrahepatic cholangiocarcinoma (ICC) cell line, RBE cells, and IDH-mutant RBE cells which produce elevated D2HG levels^3^. Carboxylic groups on D2HG may covalently modify proteins by the formation of amide bonds on lysine residues, ester bonds on serine/threonine/tyrosine residues, or thioester bonds through cysteine residues (**Fig. S1A**). We identified 41 modified sites on 23 peptides with a mass shift of 130.0266 Da, primarily O-linked (26 sites), with fewer N-linked (6) and S-linked (9) forms (**Table S1, Fig. S1B**). Our focus therefore was on *O*-2-hydroxyglutarylation of serine, threonine, and tyrosine as the O-modifications are dominant through the initial screening, but other linkage types remain to be further explored.

Our initial screening also revealed the low abundance and complexity of D2HG modifications and highlighted the need for enrichment of D2HG-modified proteins/peptides prior to LC-MS profiling. Antibody-based enrichment is effective but difficult to implement for low-abundance modifications due to the need for prior knowledge on the sites and specificity. We therefore attempted a chemical affinity enrichment based on immobilized metal ion affinity capturing for D2HG-modified peptides (**Fig. S2A**). Using polyMAC, a material originally developed for phosphopeptide enrichment^4^, we adjusted the enrichment protocol to target D2HG-modified peptides. Additionally, we used a trypsin-GluC sequential digestion strategy to minimize the interference from acidic aspartic acid (Asp) and glutamic acid (Glu) residues. As a result, the use of polyMAC enrichment in combination with sequential trypsin-GluC digestion resulted in over 10-fold enrichment compared to initial screening (**Fig. S2B, Table S2**). Further optimization of the enrichment condition indicated that increasing buffer acidity and glycolic acid (GA) concentration improved the identification of 2HG-modified peptides by protonating carboxylic groups, reducing interference from non-modified peptides. However, excessive increases in acidity and GA concentration weakened interactions between 2HG-modified peptides and IMAC, reducing enrichment efficiency (**Fig. S2C**). Analysis of 2HG modification sites revealed a serine > threonine > tyrosine distribution, similar to phosphorylation patterns^5^, suggesting potential crosstalk between these PTMs (**Fig. S2D, Table S3**).

D2HG may modify proteins through two carboxylic and one hydroxy groups (**Fig. S1A**), but we reason that the alpha carboxyl group is likely the most reactive, forming an ester bond, supported by the previously reported formation of a covalent bond by D2HG-CoA at the alpha position^6^ and the absence of the 46 Da neutral loss (H_2_CO_2_) characteristic of alpha carboxyl fragmentation^7^ in our data (**Fig. S1B**).

To validate the novel D2HG modification, we performed quantitative proteomics on H293T cells treated with exogenous D2HG. Compared to PBS only controls, D2HG-treated cells showed a dose-dependent increase in cellular D2HG concentration (**Fig. S3A**) and D2HG- modified peptides (**Fig. 1A, Table S4**). Time-dependent treatment also increased identification of D2HG-modified peptides (**Fig. S3B, Table S5**).

**Figure 1.**
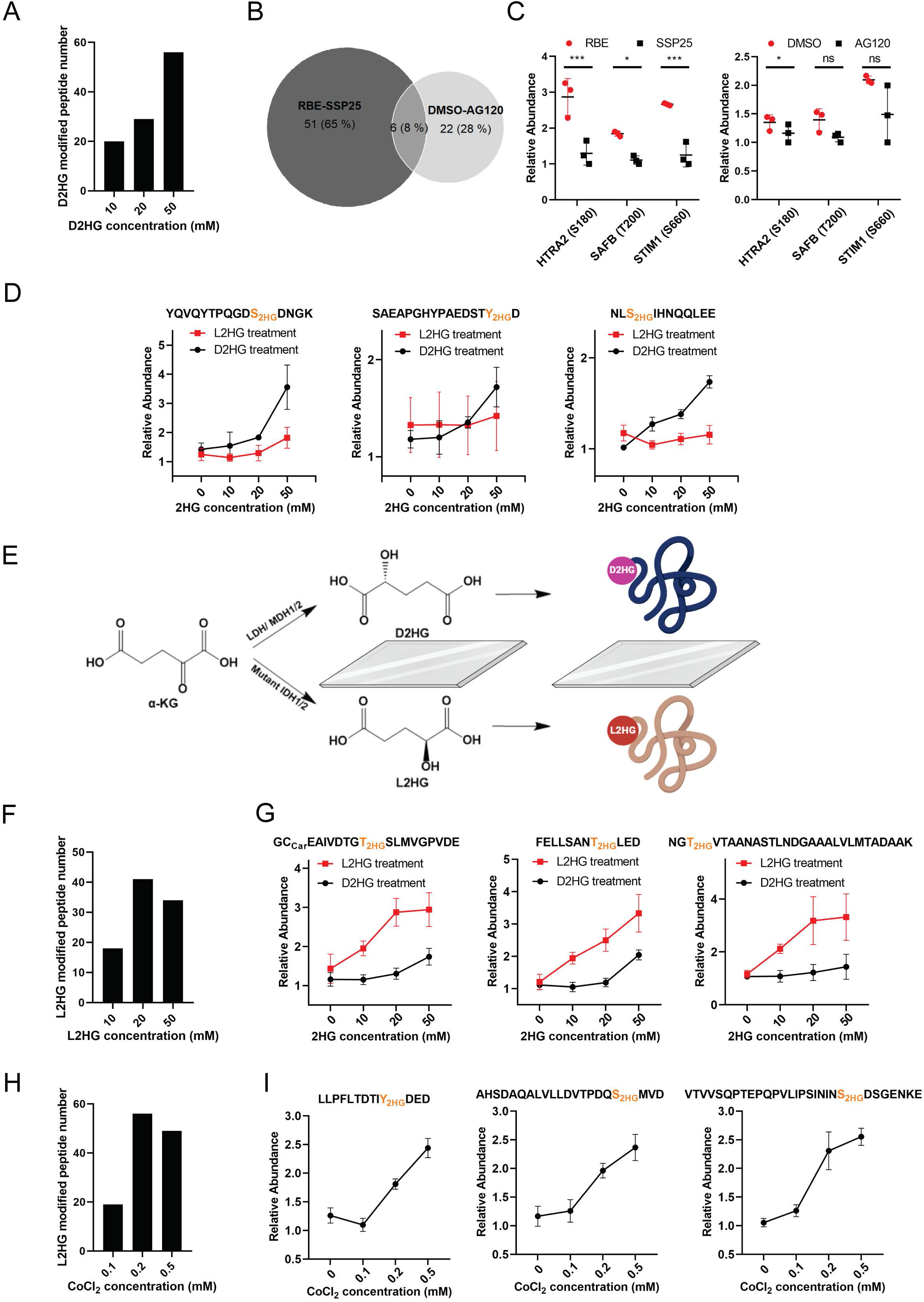
Chiral preference of D2HG and L2HG modification on proteins. (A) Identification of D2HG-modified peptides through differential proteomics. H293T cells were treated with different D2HG concentrations for 24 hours. (B) Overlap analysis of the D2HG-modified peptides identified in both the RBE-SSP25 cells and AG120 treated/ untreated RBE cells. (C) Abundance of representative D2HG modified peptides derived from RBE cells, SSP25 cells, and AG120 treated/ untreated RBE cells. (D) Abundance of representative D2HG modified peptides in H293T cells treated with a gradient concentration of D2HG and L2HG. (E) Scheme for D2HG and L2HG modification on specific proteins in a chirality behaviour. (F) Identification of L2HG-modified peptides via differential proteomics. H293T cells were treated with different L2HG concentrations for 24 hours. (G) Abundance of representative L2HG modified peptides in H293T cells treated with a gradient concentration of D2HG and L2HG. (H) Identification of L2HG-modified peptides via differential proteomics. H293T cells were treated with different CoCl_2_ concentrations for 24 hours. (I) Abundance of representative L2HG modified peptides in H293T cells treated with a gradient concentration of CoCl_2_. For differential proteomics, the peptides with a fold change > 1.2 and *P* value < 0.05 in the D2HG (or L2HG) accumulated groups compared with the non-treatment control group were considered as D2HG (or L2HG) modified peptides.

A previous study introduced AG120 that inhibits the production of D2HG by targeting mutant IDH1 **(Fig. S3C)**^8^. We examined endogenous D2HG modifications in IDH1 mutant RBE cells and AG120-treated RBE cells, identifying 79 D2HG-modified peptides by quantitative proteomics (**Fig. 1B, Table S6**). Several of these peptides showed reduced abundance following AG120 treatment (**Fig. 1C, Fig. S4**). Notably, these D2HG-modified proteins include HTRA2, SAFB, and STIM1, which are all implicated in tumor progression^9-11^, suggesting a regulatory role for D2HG in cancer.

Inspired by recent research on mirror-image biology^12^ and enantiomeric preferences of chemical probes^13^, we tested whether D2HG modifications are chiral-dependent by treating cells with D2HG and L2HG in parallel. For proteins showing dose-dependent 2HG modifications with D2HG treatment, L2HG treatment did not significantly increase modifications, indicating that the effect is primarily due to D2HG and is indeed chirality-dependent (**Fig. 1D, Table S4, Table S7**). Corresponding chromatograms and MS/MS spectra of representative peptides are shown in **Fig. S5**.

We observed some peptides with mass shifts similar to D2HG, but their levels did not correlate with exogenous D2HG treatment concentrations. We hypothesize that some of these peptides may originate from L2HG (**Fig. 1E**). To validate this, we treated cells with different concentrations of L2HG. L2HG treatment led to increases in cellular 2HG levels (**Fig. S6A**) and interestingly, parallel increase in the identification of 2HG-modified peptides (**Fig. 1F, Table S7**). Prolonged L2HG treatment produced similar effects (**Fig. S6B, Table S8**). Three representative L2HG-modified peptides, which increased significantly with L2HG compared to D2HG treatment, suggest these modifications are primarily L2HG-induced (**Fig. 1G, Fig. S7**).

It has been reported that the levels of the L-enantiomer of 2HG (L2HG) are increased by changes in the tumor microenvironment, including hypoxia and a low pH. Indeed, hypoxia induced by CoCl_2_ or low O_2_ culture, which promotes L2HG accumulation (**Fig. S6C**)^14^, increased L2HG-modified peptides (**Fig. 1H, Fig. S6D, Tables S9-S10**), with three representative peptides correlating with CoCl_2_ levels (**Fig. 1I, Fig. S8**).

To further examine MS/MS fragmentation of D2HG- or L2HG-modified peptides and minimize false positive discoveries, we noted that ester modifications, like phosphorylation, often yield neutral loss ions as the additional evidence for PTMs^15^. For D2HG and L2HG, 130.0266 Da and 148.0372 Da neutral loss ions from ester bond cleavage were observed (**Fig. S9A**). Manual inspection for the detected D2HG- or L2HG- modified peptides confirmed that most modified peptides exhibited neutral loss ions, though some at low abundance (**Fig. S9B**). Two representative MS/MS spectra display clear b/y ions and neutral loss ions (**Fig. S9C-D**), collectively validating D2HG and L2HG modifications. Comparison of 2HG-modified and unmodified peptides revealed increased retention times for modified peptides on reversed-phase columns (**Fig. S9E**). Calculated LogP values by ACD/Labs showed higher hydrophobicity in 2HG-modified serine (−0.97 ± 0.45) than in unmodified serine (−1.58 ± 0.33), which aligns with observed retention shifts and could serve as an additional indicator of 2HG modifications.

To validate and enhance specificity, we used isotopic metabolic flux analysis by incubating cells with isotopically labelled D/L2HG-d4 to directly track potential incorporation of isotopic 2HG tag. The isotopic metabolic flux experiment allowed us to identify 124 d4-2HG-modified peptides (**Table S11**). MS/MS spectra of heavy-labelled and unlabelled 2HG-modified peptides showed identical fragmentation with a +4 Da shift, confirming endogenous 2HG modification structure (**Fig. S10**).

Overall, 56 high-confidence D2HG modifications were identified in D2HG-accumulated cells, with significant overlap between IDH mutant RBE cells and H293T cells; while 130 high-confidence L2HG modifications were identified in L2HG accumulated cells (detected at least twice with abundance correlating to cellular D2HG or L2HG levels, **Tables S12-S13**). D2HG- and L2HG-modified proteins showed distinct patterns, with 33 D2HG-specific and 87 L2HG-specific proteins (**Fig. S11A**). Gene Ontology (GO) analysis revealed distinct functional roles: D2HG-modified proteins were enriched in ATP binding and chromatin remodelling, while L2HG-modified proteins were associated with RNA binding and oxidative stress response (**Fig. S11B-E**). These functions align with the respective roles of D2HG and L2HG in genome regulation and hypoxia response^16^. The analysis (**Fig. S11B**) also revealed 2HG-modified proteins associated with kinase signaling. To investigate the effect of 2HG modifications on kinases, we manually examined all 2HG-modified kinases in our data. As representative, 2HG modification was observed at Ste20-like serine/threonine kinase (SLK) S719, with levels increasing following exogenous L2HG treatment, while D2HG had little effect (**Fig. 2A**). CoCl_2_ treatment in H293T cells also elevated L2HG-modified SLK, correlating with cellular L2HG levels (**Fig. 1I right, Fig. S6C**). This modification was confirmed with the synthetic 2HG-modified peptide with essentially identical retention time and MS/MS spectra (**Fig. 2B, 2C**). Furthermore, SLK is known to phosphorylate PLK1 at T210^17^, and phosphoproteomics showed reduced PLK1 T210 phosphorylation in CoCl_2_- treated cells, suggesting L2HG modification potentially inhibits SLK activity (**Fig. 2D**).

**Figure 2.**
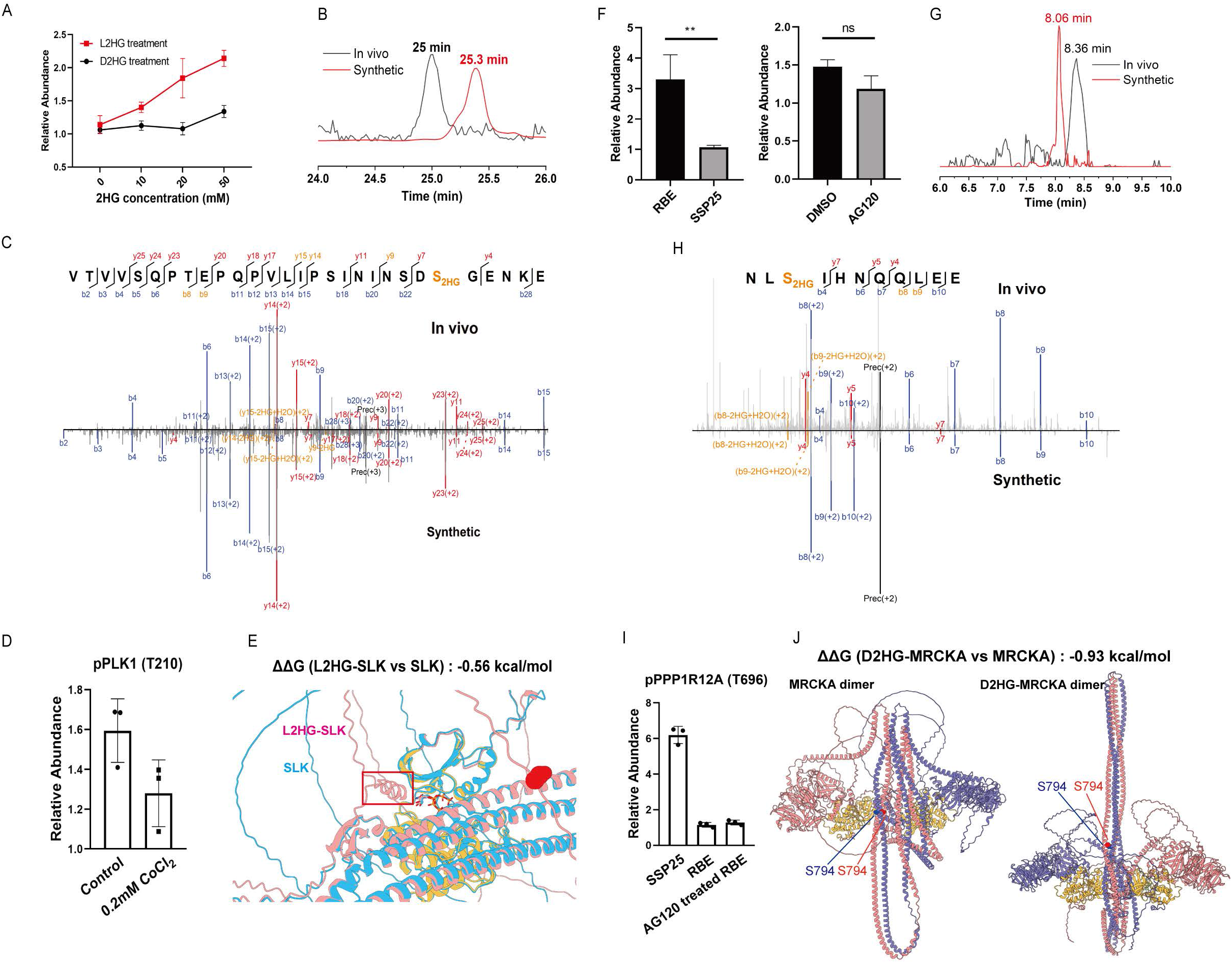
Kinase SLK modified with L2HG and MRCKA modified with D2HG possess decreased activity. (A) Quantification of L2HG modified SLK (S719) in H293T cells treated with varying concentration of D2HG and L2HG. (B-C) Comparison of EICs and MS/MS spectra between in-vivo detected peptide and synthetic standard for SLK (S719). (D) Abundance of phosphorylated PLK1 (T200) in H293T cells treated with or without 0.2 mM CoCl_2_. (E) Structures of the SLK with or without L2HG modification. Light blue represents SLK, pink represents L2HG- SLK, and yellow represents active pocket. (F) Quantification of D2HG modified MRCKA (S794) in RBE cells, SSP25 cells, and AG120-treated RBE cells. (G-H) Comparison of EICs and MS/MS spectra between in-vivo detected peptide and synthetic standard for MRCKA (S794). (I) Abundance of phosphorylated PPP1R12A (T696) in RBE cells, SSP25 cells, and AG120-treated RBE cells. (J) Structures of the MRCKA dimer with or without D2HG modification. Yellow represents active pocket.

Similarly, we observed increased D2HG modification at MRCKA (CDC42BPA) S794 after D2HG treatment in H293T cells, while L2HG treatment had less effect (**Fig. 1D right**). The modification also correlated with endogenous D2HG levels in ICC cells of different genotypes (**Fig. 2F**). We further confirmed the modification with the synthetic 2HG-modified peptide (**Fig. 2G, 2H**). MRCKA, which phosphorylates PPP1R12A at T696^18^, showed decreased phosphorylation in D2HG-accumulated RBE cells compared to wild-type SSP25 cells. AG120, a mutant IDH1 inhibitor, partially reversed this effect (**Fig. 2I**). These results suggest that D2HG modification inhibits MRCKA activity, consistent with reports linking decreased MRCKA activity to slowed cell growth^19^, potentially explaining the reduced growth rate in IDH-mutant cells^20^.

To explore the mechanisms underlying reduced kinase activity with 2HG modifications, we modeled MRCKA and SLK structures using AlphaFold 3. Mimicking acidic 2HG modification via mutagenesis strategy^21^ revealed that 2HG modification of SLK (S719) created a barrier at the active site pocket, potentially hindering substrate access (**Fig. 2E**). Free energy changes (ΔΔG = -0.56 kcal/mol) further supported SLK destabilization upon S719 modification. Similarly, MRCKA’s dimerization, essential for activation^22^, was destabilized. The modification (S794), located at the coiled-coil domain responsible for dimer formation (**Fig. 2J**), likely induced electrostatic repulsion, reducing kinase activity. Free energy calculations supported this hypothesis (ΔΔG = -0.93 kcal/mol).

In summary, we report here the novel covalent protein modifications by oncometabolite D2HG in ICC cells with IDH1/2 mutation. We further discovered chiral dependent protein modifications by D2HG and L2HG. These novel covalent modifications were confirmed through exogenous treatment, isotopic metabolic flux analysis, neutral loss in tandem mass spectrometry, and synthetic peptide authentication. Our findings highlight D2HG and L2HG modifications as potential regulators in cancer biology and underscore the need for further study to elucidate their functional significance and mechanistic roles.

## Online content

Any methods, additional references, Nature Research reporting summaries, source data, extended data, supplementary information, acknowledgements, peer review information; details of author contributions and competing interests; and statements of data are available at http://doi.org/XXX.

## Data Availability

The proteomics data have been deposited to the ProteomeXchange Consortium via the jPOST partner repository with the dataset identifier PXD059758. All other data supporting the findings of this study are available from the corresponding author upon request.

## Acknowledgments

This project was partially supported by NIH grants 4R01AG064250 (to W.A.T.), 2RO1CA069202 (to Z.Y.Z.), 1R35GM138002 (to E.I.P.), and P30 CA023168 (to Purdue Institute for Cancer Research), and by NSF grant 2404098-CHE (to W.A.T.).

## Author contributions

Z.Z. and W.A.T. designed experiments. Z.Z. performed the MS experiments and analyzed the data. Y.K.L. and Z.J.L. contributed to sample preparation. M.J.W. and N.B. provided the ICC cell lines and other resources on cellular experiments. C.N.E., Z.H.Q., Z.Y.Z. and E.I.P. contributed to 2HG modified peptide standard synthesis and characterization. Z.Z. and W.A.T. wrote the paper with input from all authors. All authors reviewed and approved the manuscript.

