## Supplementary Materials for "Discovery of Chirally-dependent Protein O-2-Hydroxyglutarylation by D2HG and L2HG"

#### Methods

#### Reagents

Sequencing grade trypsin and Glu-C was obtained from Thermo Fisher (Waltham, MA) and Sigma (St. Louis, MO) respectively. polyMAC was obtained from Tymora (West lafayette, IN). Isotopic labeled d4 sodium (RS)-2-Hydroxyglutarate (2,3,3-$D_3$; OD, 98%) was purchased from Cambridge Isotope Laboratories (Tewksbury, MA), which was called as d4 D/L 2HG hereafter. Other reagent was obtained from Sigma (St. Louis, MO) unless otherwise noted.

#### Chemical synthesis of D/L 2HG modified peptides

##### (1) Synthesis of diethyl 2-hydroxypentanedioate (S1)

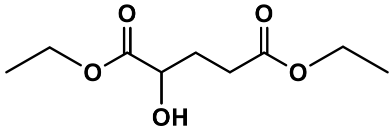


**S1**

1 eq. (1g, 5.208 mmol) of 2HG were weighted into a 250mL flask, followed by the addition of 40mL ethanol and 0.3eq. (0.4mL) of $H_2SO4$. The reaction was refluxed at 70°C overnight. The solvent was evaporated, followed by the addition of 20mL $H_2O$ and neutralization using 1N KOH. The aqueous layer was extracted 3 times with 50mL of EtOAc and dried by $Na_2SO4$ and filtered. The filtrate was concentrated in vacuo. The crude was directly used for next step synthesis.

##### (2) Synthesis of diethyl 2-(benzyloxy) pentanedioate (S2)

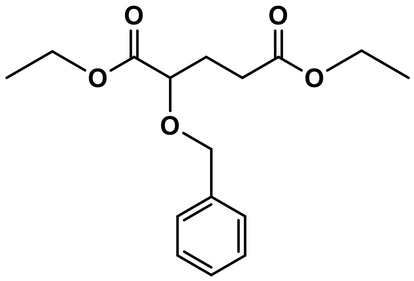

**S2**

Add dropwise a solution of Compound S1 (0.81g, 4mmol) in DMF (2mL) to a suspension of NaH (60% in mineral oil, 0.32g, 8mmol) in dry DMF (8mL) at 0°C. Stir the reaction for 30 min at 0°C. Add BnBr (0.953mL, 8mmol) to the mixture. Continue the stirring for 40 min. Stir the mixture at 0°C for 2h. Add TEA (1.13mL, 8mmol) and stirred for another 1h. Quench the reaction by addition of $H_2O$ (50mL). Extract the reaction with $Et_2O$ (5*8mL). Wash the extracts with brine and dry by $Na_2SO4$. Dry the solution and purify the product by column chromatography (petroleum ether : EtOAc = 10:1). MS (ESI): m/z calcd for $C_{16}H_{23}O_5^+$ $[M+H]^+$: 295.1; found: 295.1.

**(3) Synthesis of 2-(benzyloxy) pentanedioic acid (S3)**

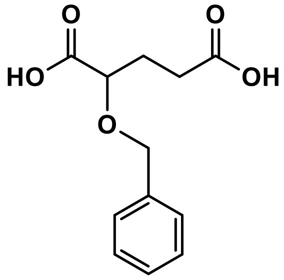

**S3**

To a solution of S2 (0.33g, 1.1224mmol) in 2.37mL EtOH at 0°C was added a solution of KOH (0.25g, 4.49mmol) in 2.37mL $H_2O$. The reaction was allowed to warm to RT and stirred overnight. After concentrating the solution in vacuo to remove EtOH, the remained aqueous phase was adjusted to pH 1-2 by HCl. Then dry the solution in vacuum. MS (ESI): m/z calcd for $C_{12}H_{13}O_5^-$ [M-H]$^-$: 237.1; found: 237.1.

**(4) Synthesis of 2,5-bis(benzyloxy)-5-oxopentanoic acid (S4)**

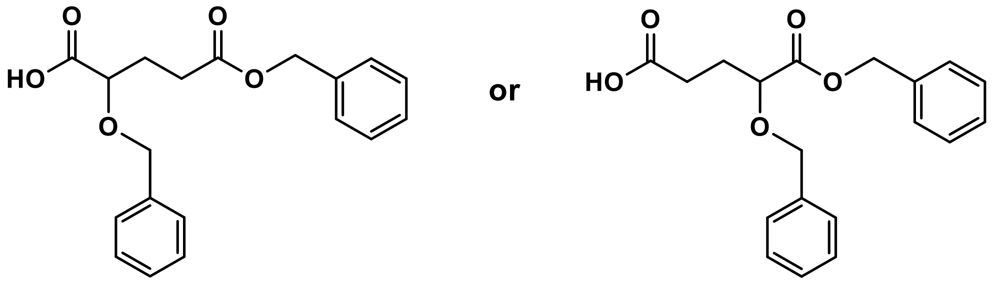


**S4**

Oxalyl chloride (1.36mmol, 0.17g, 116ul, 1.2eq) was added to a stirring solution of Compound S3 (0.27g, 1.1345mmol, 1.0eq) in DCM (1mL) at 0°C, followed by adding two drops of DMF. The reaction was allowed to react at RT for 3 h, followed by drying in vacuum. The crude was dissolved in 1mL DCM. The resulting solution was added dropwise to a solution of BnOH (1.3614mmol, 147mg, 1.2eq) and TEA (114.8mg, 1.1345mmol, 1eq) in 1mL THF at 0°C. The reaction was stirred for 1h at 0°C and for overnight at RT. The resulting solution was quenched by 0.1ml $H_2O$, followed by drying and purifying by C18-HPLC (5%-90% MeOH in 20 min). MS (ESI): m/z calcd for $C_{19}H_{19}O_5^-$ [M-H]$^-$: 327.1; found: 327.1.

**(5) Synthesis of N-(((9H-fluoren-9-yl)methoxy)carbonyl)-O-(2,5-bis(benzyloxy)-5-oxopentanoyl)-L-serine [Fmoc-Ser(2HG)-OH, S5]**

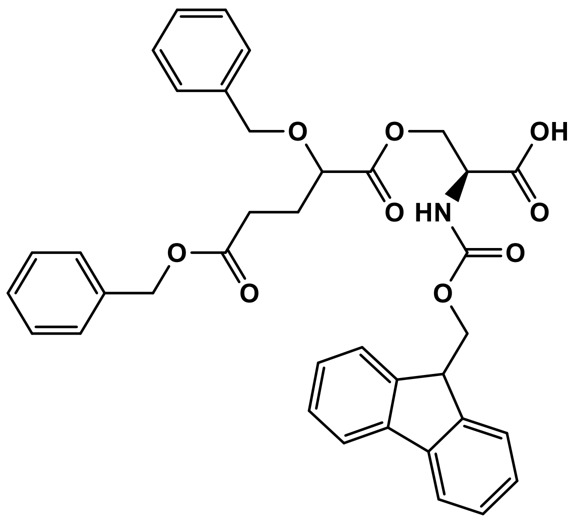

**S5**

Oxalyl chloride (87mg, 0.6860mmol, 1.5eq) was added to a stirring solution of Compound S4 (150mg, 0.4573mmol, 1.0eq) in DCM (1mL) at 0°C, followed by adding two drops of DMF. The reaction was allowed to react at RT for 4 h, followed by drying in vacuum. The crude was dissolved in 1mL DCM. The resulting solution was added dropwise to a solution of Fmoc-Ser-OH (224.3mg, 0.686mmol, 1.5eq) and TEA (92.55mg, 0.9146mmol, 2eq) in 1mL DCM at 0°C. The reaction was stirred for 1h at 0°C and for 1h at RT. The resulting solution was dried and the isomers were purified by C18-HPLC (5%-90% MeOH in 30 min). HRMS (ESI): m/z calcd for $C_{37}H_{36}NO_9^+$ $[M+H]^+$: 638.239; found: 638.235. 1H-NMR (300 MHz, $CDCl_3$) δ 7.75 (d, 4H, CH=C), 7.60 (d, 4H, CH=C), 7.26-7.40 (m, 10H, CH=C), 5.80 (d, 1H, CONH), 5.3 (s, 2H, $COOCH_2$), 5.2 (m, 2H, $COOCH_2$), 5.05 (d, 1H, COCH), 4.30-4.46 (m, 6H, $OCH_2$), 2.45 (m, 2H, $COCH_2$), 2.10 (m, 2H, $COCH_2CH_2$).

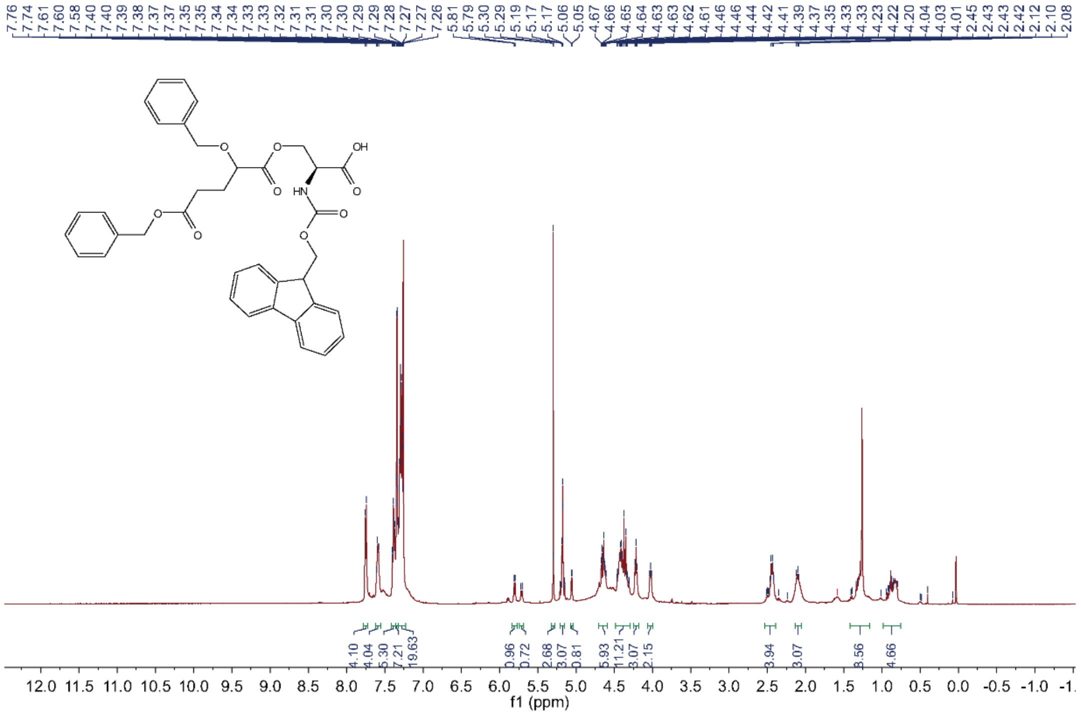


**(6) Peptide standard synthesis**

Compound S5 served as the protected amino acid for peptide synthesis. Peptide standards containing the 2HG modification were synthesized using a standard Fmoc-based solid-phase peptide synthesis workflow[1].

**Cell Culture and treatment**

ICC cell lines with different genotypes were generously provided by Dr. Nabeel Bardeesy (Massachusetts General Hospital, Harvard Medical School, Boston, Massachusetts). Cells were cultured in RPMI medium (Thermo Fisher Scientific Inc., Waltham, MA) with 10% FBS at 37°C in 5% $CO_2$. For hypoxic conditions, cells were treated with varying concentrations of the hypoxia inducer $CoCl_2$ (0, 100, 200, and 500 μM) for 24 hours or grown in a specialized, humidified chamber equilibrated with 1% oxygen, 94% nitrogen, and 5% carbon dioxide for the indicated duration.

For isotopic metabolic flux experiments, cells were washed once with warm PBS and replenished with fresh RPMI with 10% FBS before labeling by d4 D/L 2HG. D4 D/L 2HG (25 mM) was added into the media and cells were collected after 24h.

**Cell harvest and lysis**

Cells were harvested when reaching about 80% confluent. Cells were washed with cold PBS for two times and scraped from dish and transferred to a microcentrifuge tube. After that, cells were resuspended in lysis buffer (8M urea in 50mM Tris-HCl buffer, pH 8.0) with 1 tablet of EDTA-free protease inhibitor Cocktail, and then sonicated on ice for three 1 min rounds at 15% amplitude. The lysates were then centrifuged at 12,000 rpm, 4°C for 20 min. The supernatant was transferred to a new tube. Total protein concentrations were determined by bicinchoninic acid (BCA) protein assay kit (Thermo Scientific Pierce, Rockford, IL).

Cells were harvested at approximately 80% confluence, washed twice with cold PBS, and scraped from the dish into microcentrifuge tubes. The cells were then resuspended in lysis buffer (8 M urea in 50 mM Tris-HCl, pH 8.0) containing an EDTA-free protease inhibitor tablet. The suspension was sonicated on ice for three 1-minute rounds at 15% amplitude. Lysates were centrifuged at 12,000 rpm for 20 minutes at 4°C, and the supernatant was transferred to a new tube. Total protein concentrations were measured using a bicinchoninic acid (BCA) protein assay kit (Thermo Scientific Pierce, Rockford, IL).

**Trypsin/Glu-C sequential digestion**

1M DTT stock solution was added into cell lysates to a final concentration of 10mM. The obtained solution was vortexed and incubated at 30°C for 45 min. Then 0.6M CAA stock solution was added to a final concentration of 30mM, followed by vortexed and incubated at RT in the dark for 30 min. Afterwards, another 1M DTT was added to a final concentration of 10mM, followed by vortexed and incubated at RT for 10 min. The obtained solution was diluted with 50mM ammonium bicarbonate to make urea concentration less than 1M. Trypsin (enzyme: sample = 1:50) was added into the solution and incubated at 37° C for 16 h. Then the same volume of phosphate buffer (pH 7.5) and Glu-C (enzyme: sample = 1:50) and was added into the solution and incubated at 37°C for 8h. The reaction was quenched by acidification with 1% TFA, followed by desalting with C18 spin columns. The obtained solution was lyophilized for further use.

**Enrichment of 2-HG modified peptides by polyMAC**

The 2HG peptides were enriched using polyMAC beads. Firstly, 50 μL of polyMAC beads (Tymora, West lafayette, IN) were rotated for 1 min and quickly spun down. The supernatants were removed, and 200 μL of Loading buffer (0.01% TFA, 20mM glycolic acid (GA), 50% ACN) was added to the tubes, followed by rotation, centrifugation, and removal of supernatants.

Next, 500 μL of loading buffer was added to the polyMAC, and the beads were split into three tubes (Tube I: 200 μL; Tube II: 200 μL; Tube III: 100 μL). Enrichment was then performed using the sequential enrichment method.

100 µg of peptides dissolved in 500 µL of loading buffer was added to Tube I, followed by incubation at room temperature with shaking for 30 min. The polyMAC beads were then spun at 2,000g for 1 min, and the supernatant was transferred to Tube II. This was incubated at room temperature with shaking for 20 min, followed by spinning at 2,000g for 1 min and transferring the supernatant to Tube III. Finally, the mixture was incubated at room temperature with shaking for 10 min.

After that, the polyMAC beads in tube III were spun down at 2000g for 1 min and the supernatants were discarded. The three tubes of polyMAC were combined by using 200 uL of Washing buffer A (0.1% TFA in 50% ACN), followed by spinning down at 2000g for 1 min and discarding the supernatants. The polyMAC beads were then washed with 200 µL of Washing buffer A (0.1% TFA in 50% ACN) four times and 200 µL of Washing buffer B (80% ACN) one time. After centrifugation at 2000g for 1 min and discarding the supernatants, the peptides were eluted from polyMAC beads with 150 µL of Elution buffer (400mM $NH_4OH$ in 50% ACN) by shaking at 1,100 rpm for 30 min at RT. The elution step was repeated once. The obtained eluted solution were collected and dried down in a SpeedVac and stored at -80°C.

**Phosphopeptides enrichment**

The phosphopeptides were enriched using polyMAC beads. Firstly, 100 µg dried tryptic peptides were dissolved into 200 µl of Loading buffer (100mM glycolic acid, 1% TFA, 50% ACN) in a non stick microfuge tube. Then 50 µl of polyMAC reagent was added and shake for 15 min. After that, pipette the mixture up-and-down a few times and transfer the whole solution to the spin column and spin down at 5,000 rpm for 30 seconds to remove the unbonded peptides. Then the polyMAC beads were sequentially washed with Loading buffer, Washing buffer I (25 mM glycolic acid, 0.2% TFA, 80% ACN) and Washing buffer II (80% ACN) for three times. Finally, the attached phosphopeptides were eluted from the polyMAC beads by 50 μl Elution buffer (400 mM $NH_4OH$, 50% ACN) for 2 times, followed by drying the samples in SpeedVac.

**LC-MS analysis**

Samples were measured on a timsTOF pro2 (Bruker Daltonics Gmbh) with a reversed phase Evosep One (Evosep). For peptide separation, we used the standard SPD30 or SPD60 methods. The mass spectrometer was operated in a data-dependent acquisition parallel accumulation-serial fragmentation (PASEF) mode with ten PASEF scans per topN acquisition cycle. The accumulation and ramp time for the dual TIMS analyzer were set to 100 ms as a duty cycle. Full MS data were acquired in a mass range of 100–1700 m/z and mobility range of 0.6–1.6 V $s^{-1}$ $cm^{-2}$ 1/*K0*. The collision energy was ramped as a function of increasing mobility starting from 20 eV at 0.6 V $s^{-1}$ $cm^{-2}$ to 60 eV at 1.6 V $s^{-1}$ $cm^{-2}$.

**Data analysis**

The raw files were processed using the FragPipe software 21.0, a robust software suite for DDA analysis. The data was searched against a reverse concatenated, nonredundant variant of the Human UniProt database (UniProt_Human_reviewed_26-03-2024.fasta) by the engine MSFragger (v4.0). A mass tolerance of 20 ppm was allowed for precursor ions and 20 ppm for fragment ions. Cysteine carbamidomethylation (+57.0215 Da) was chosen as static modification. 130.0266 Da (for 2HG modification) was set as variable modifications on serine, threonine and tyrosine. 15.9949 Da was set as variable modification on methionine. Trypsin, GluC or trypsin-GluC was set as the protease, allowing a maximum missed cleavage of two, two, or three respectively. The search results were statistically validated using Philosopher (v5.1.0), and PTMProphet was employed to compute PTM site localization probabilities. IonQuant (v1.10.12) was used for MS1 precursor intensity quantification, applying a minimum PTM site probability threshold of 0.75.

Peptide-spectrum matches (PSMs) were filtered to 1% FDR using a target-decoy database approach. Missing values were imputed using the minimum values in each dataset[2]. To identify high-confidence 2HG-modified peptides, we applied two criteria: peptides with fold change > 1.2 and $P < 0.05$ in D2HG or L2HG-accumulated groups were whitelisted, while those with fold change < 0.833 and $P < 0.05$ were blacklisted and excluded. Peptides identified only once as modified were also excluded.

MS/MS spectra of 2HG-modified peptides were manually inspected, requiring detection of at least two b/y ions containing the 2HG site. For quantification of 2HG modification and phosphorylation sites, we manually examined the extracted ion chromatograms (EICs) of modified peptides and integrated their peak areas.

**Functional annotation and statistical analyses**

The 2HG modified proteins were subjected to GO functional annotation including molecule function and biological process by DAVID Bioinformatics Resources 6.8 (https://david.ncifcrf.gov/). For comparison between two groups, datasets were analyzed by a two-tailed Student's t-test.

#### Prediction of 2HG modified protein models

MRCKA and SLK were selected as representative proteins for modelling with or without 2HG modifications. Protein structures were generated using AlphaFold 3[3]. To mimic acidic 2HG modifications, serine residues at the modification sites were substituted with asparagine using conventional mutagen esis strategy[4, 5]. The thermodynamic stability of the proteins was assessed via changes in folding free energy (ΔΔG), calculated using SAAFEC-SEQ[6].

### Supplementary Figures

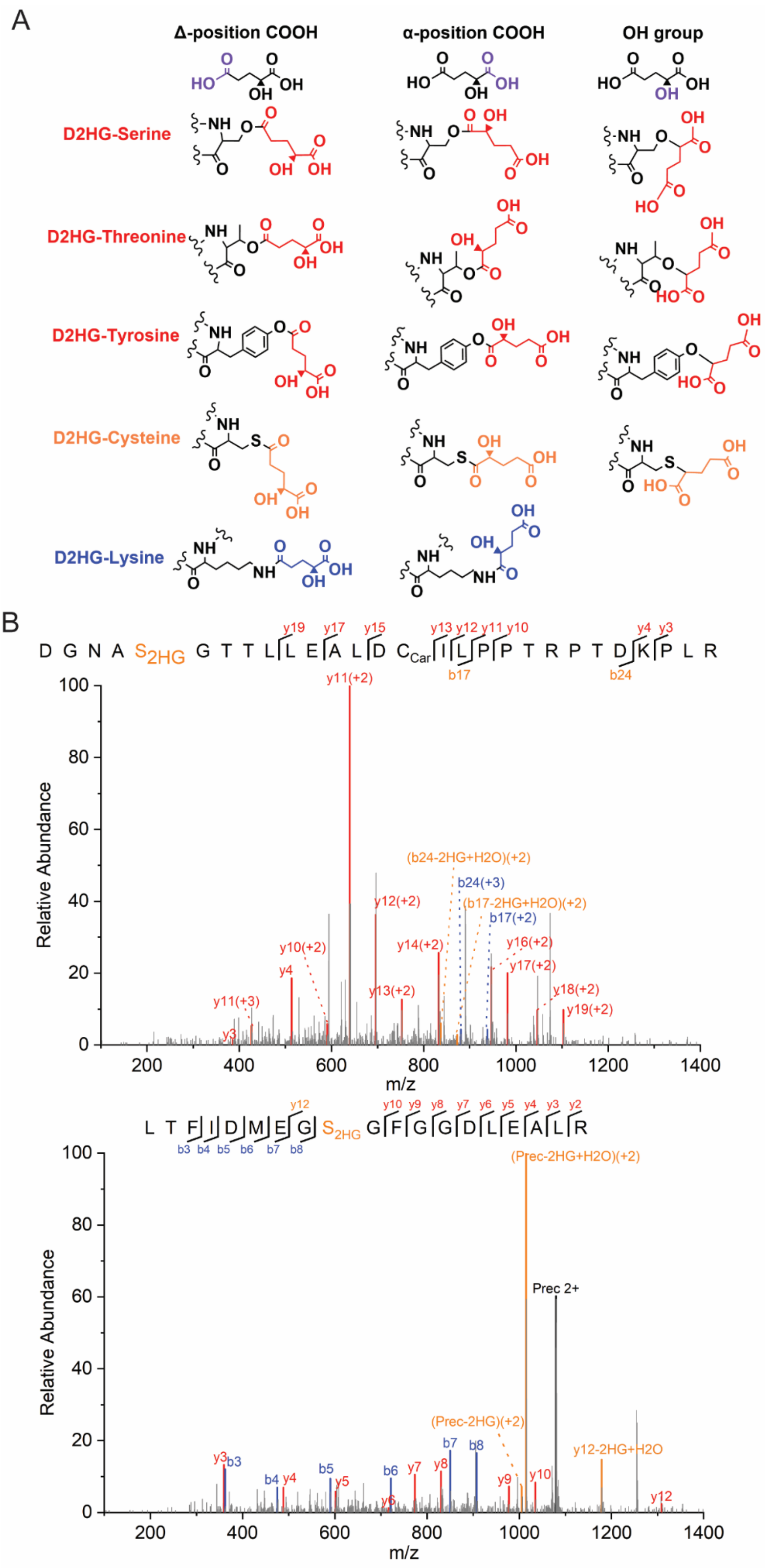

A
Δ-position COOH
α-position COOH
OH group
D2HG-Serine
D2HG-Threonine
D2HG-Tyrosine
D2HG-Cysteine
D2HG-Lysine
B
D G N A S2HG G T T L L E A L D CCar I L P P T R P T D K P L R
Relative Abundance
m/z
y11(+2)
(b24-2HG+H2O)(+2)
b24(+3)
(b17-2HG+H2O)(+2)
b17(+2)
y12(+2)
y14(+2)
y16(+2)
y17(+2)
y18(+2)
y19(+2)
y10(+2)
y13(+2)
y11(+3)
L T F I D M E G S2HG G F G G D L E A L R
(Prec-2HG+H2O)(+2)
Prec 2+
(Prec-2HG)(+2)
y12-2HG+H2O

**Fig S1. Exploration of potential D2HG modifications.** (A) Possible structure of D2HG modification. (B) MS/MS spectra of representative peptides with 2HG modifications. The b ion marked as blue refers to the N-terminal parts of the peptide, the y ion marked as red refers to the C-terminal parts of the peptide, and the ion with neutral loss of 2HG were marked as orange colour.

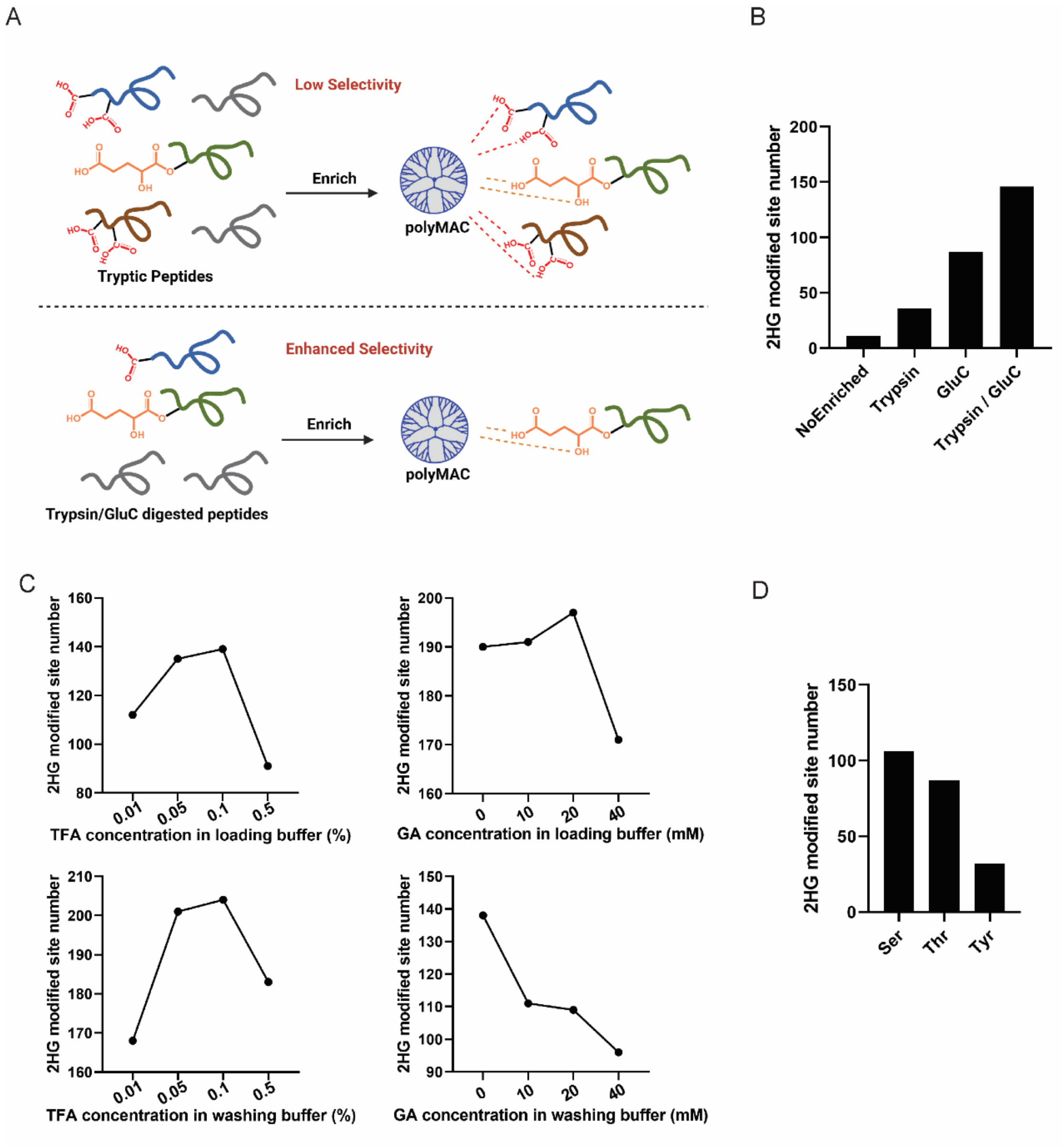


**Fig. S2. Investigation of polyMAC enrichment for 2HG modification.** (A) Schematic representation of the polyMAC enrichment strategy for D2HG modifications. (B) Comparison of the number of identified 2HG modification sites using different enrichment approaches. (C) Method optimization for the polyMAC enrichment of 2HG-modified peptides digested by trypsin-GluC. (D) Site distribution analysis for identified 2HG modified peptides.

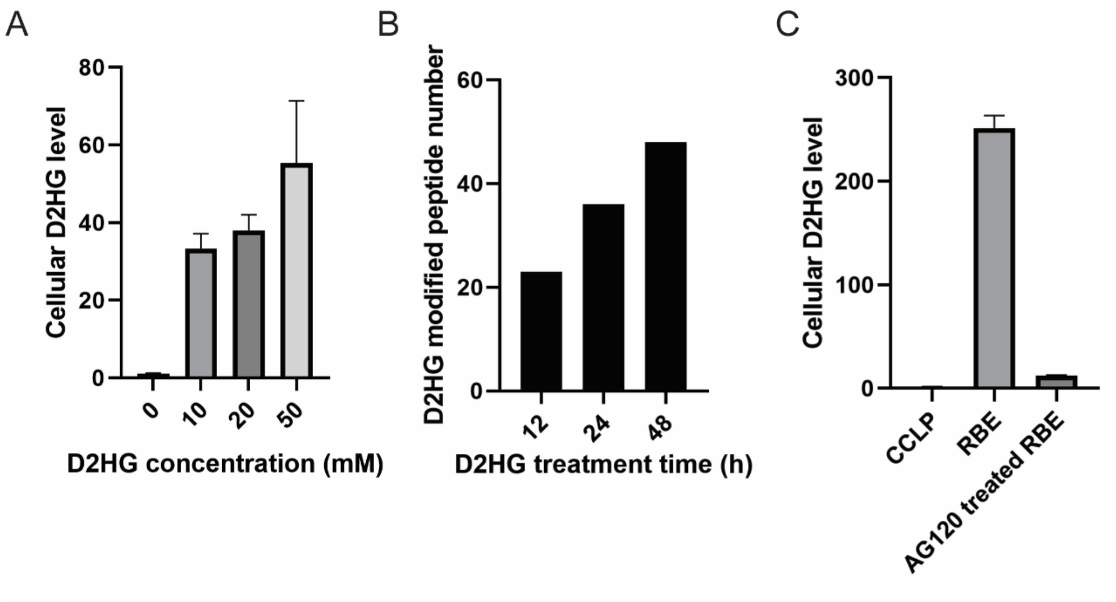


**Fig. S3. Recognition of D2HG modified peptides.** (A) LC-MS determination of D2HG concentrations in H293T cells with different concentration of D2HG treatment. (B) H293T cells treated with 20 mM D2HG over different time points. The peptides with a fold change > 1.2 and *P* value < 0.05 in the D2HG treatment groups compared with the non-treatment control group were considered as D2HG modified peptides. (C) LC-MS quantification of D2HG concentrations in wild-type SSP25 cells, IDH1 mutant RBE cells, and AG120-treated RBE cells.

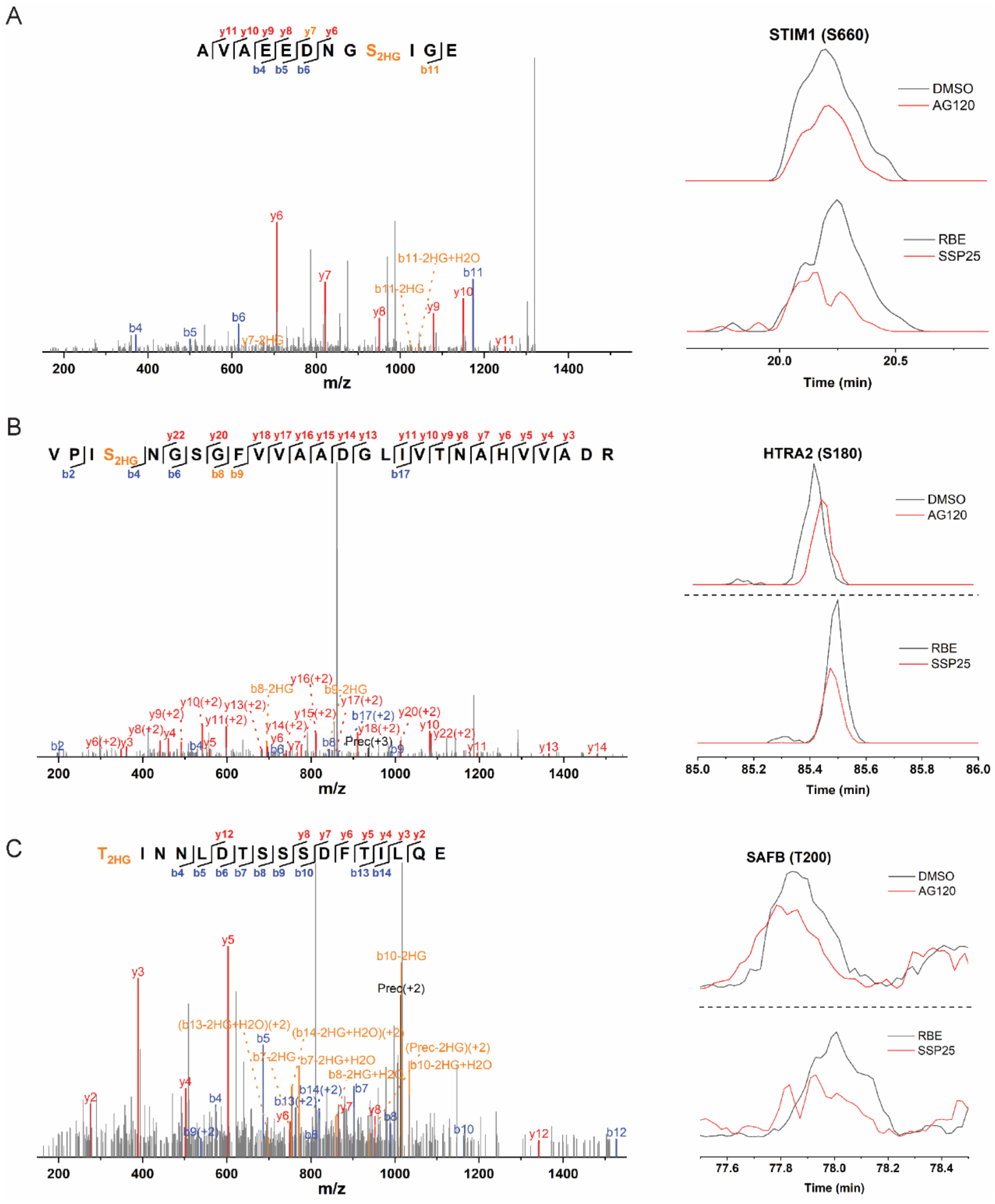


**Fig. S4.** Extracted ion chromatograms (EICs) and MS/MS spectra of representative D2HG modified peptides derived from IDH1 mutant RBE cells, wild type SSP25 cells, and RBE cells treated with or without AG120 (a mutant IDH1 inhibitor). The neutral loss ions are highlighted in orange, the b ions are shown in blue and the y ions are shown in red.

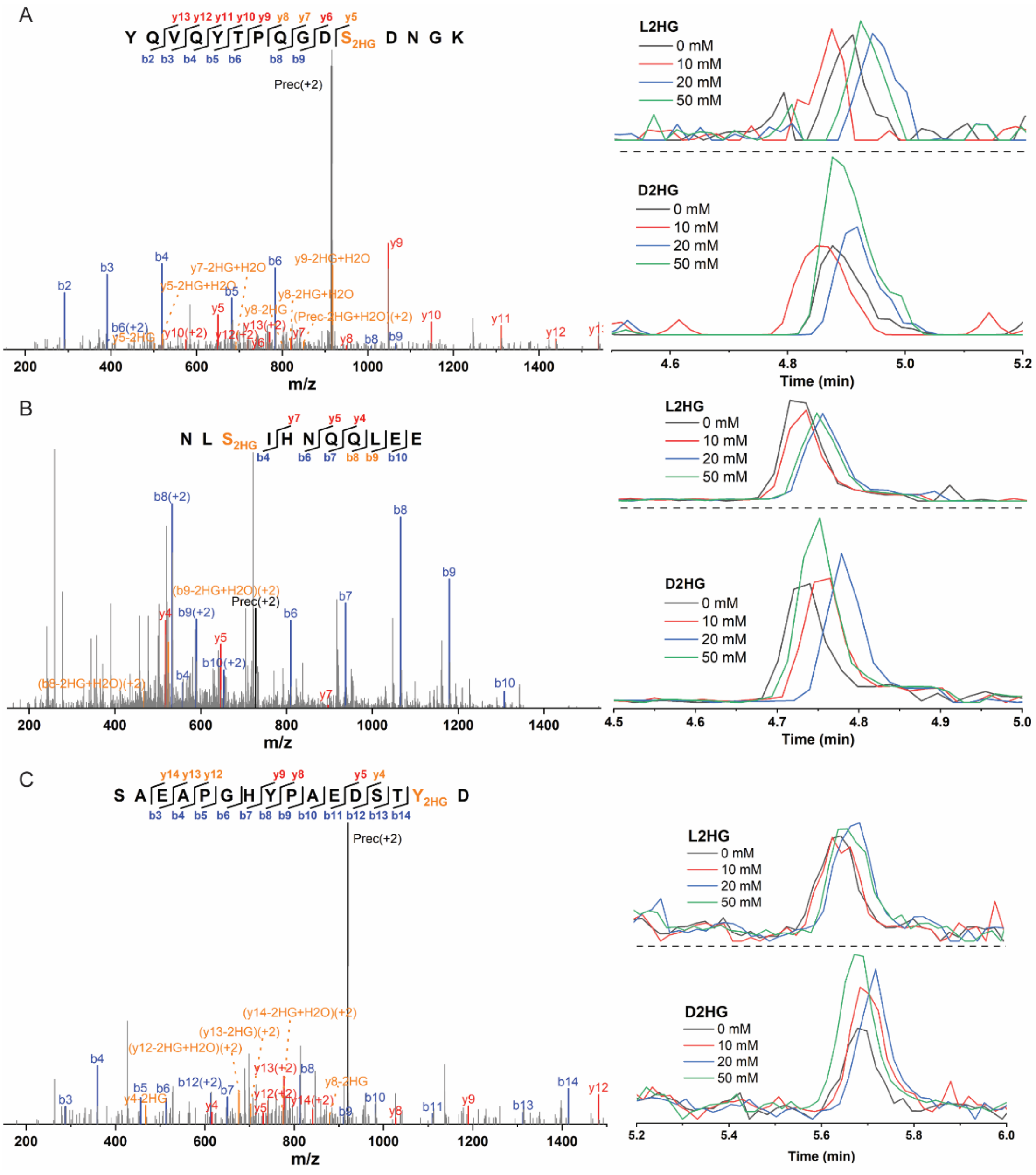


**Fig. S5.** MS/MS spectra and extracted ion chromatograms (EICs) of representative D2HG modified peptides detected in H293T cells treated with exogenous D2HG and L2HG. The neutral loss ions are highlighted in orange, the b ions are shown in blue and the y ions are shown in red.

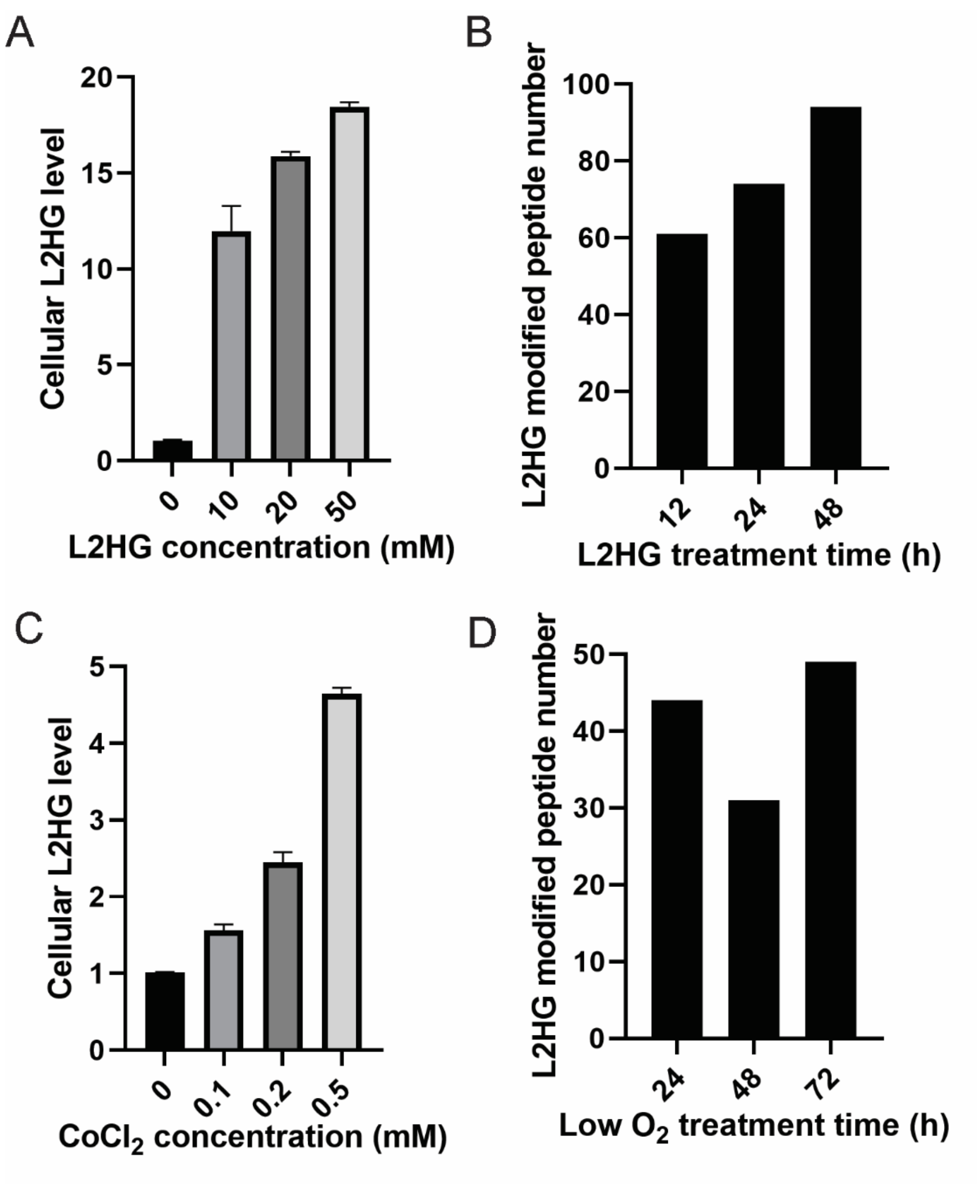


**Fig. S6. Recognition of L2HG modified peptides.** (A, C) LC-MS quantification of L2HG concentrations: (A) H293T cells treated with various concentrations of L2HG, and (C) H293T cells treated with $CoCl_2$ at different concentrations. (B, D) Identification of L2HG-modified peptides via differential proteomics: (B) H293T cells treated with 20 mM L2HG over varying time points; (D) H293T cells exposed to a low $O_2$ environment for different durations. Peptides with a fold change > 1.2 and P value < 0.05 in the treatment groups, compared to the untreated control group, were classified as L2HG-modified peptides.

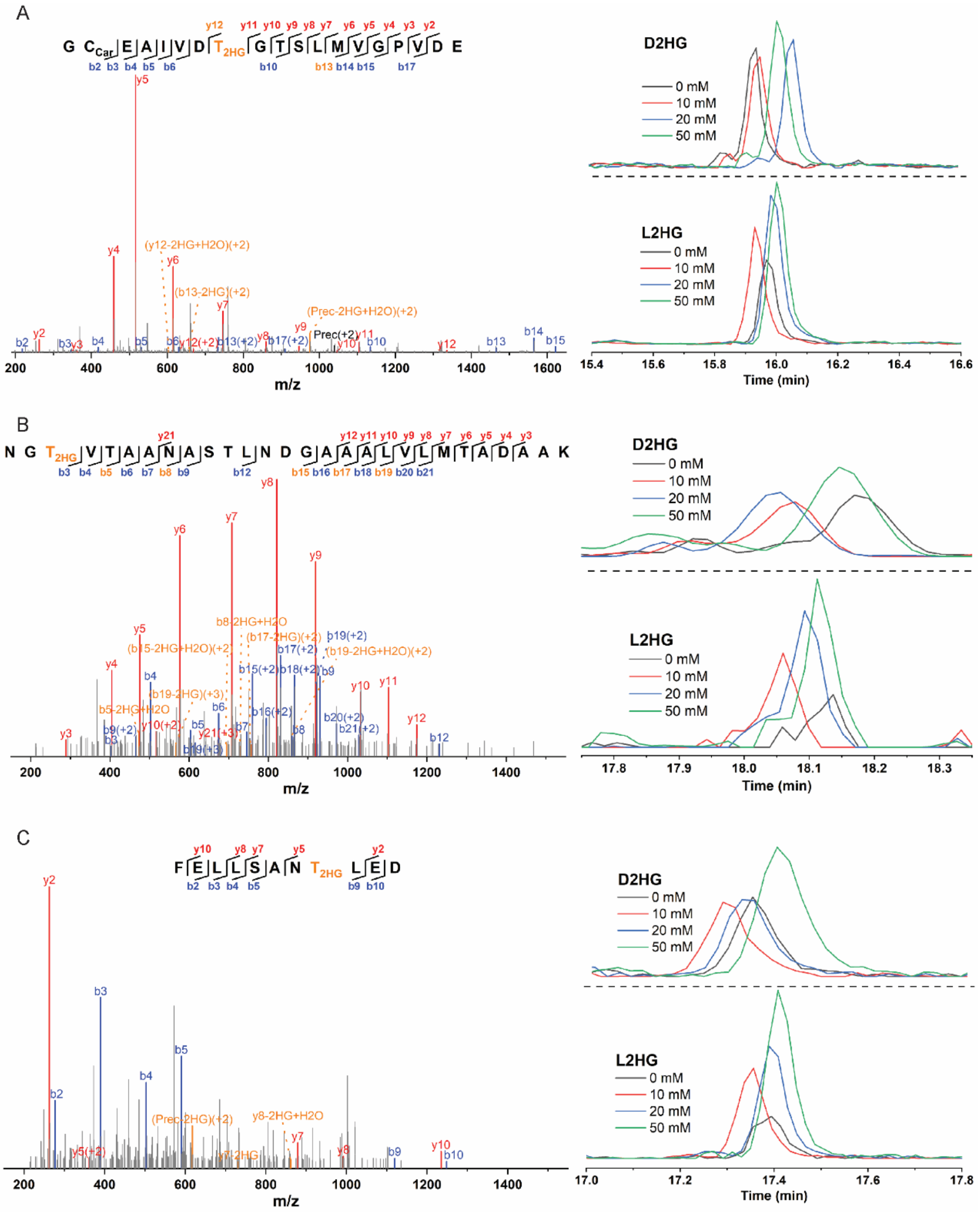


**Fig. S7** MS/MS spectra and EICs of representative L2HG modified peptides detected in H293T cells treated with exogenous D2HG and L2HG. The neutral loss ions are highlighted in orange, the b ions are shown in blue and the y ions are shown in red.

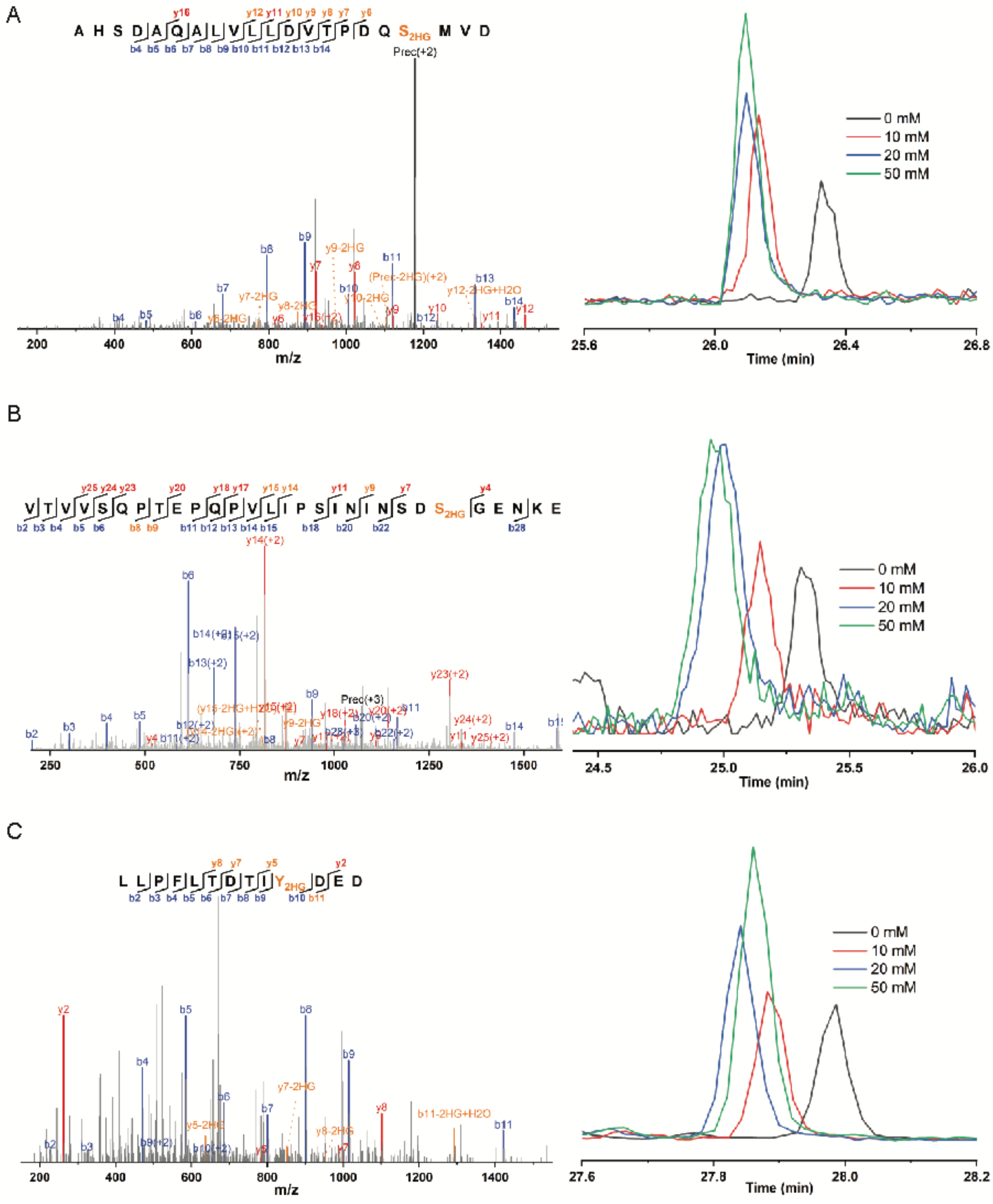


**Fig. S8.** MS/MS spectra and EICs of representative L2HG modified peptides detected in H293T cells treated with $CoCl_2$. The neutral loss ions are highlighted in orange, the b ions are shown in blue and the y ions are shown in red.

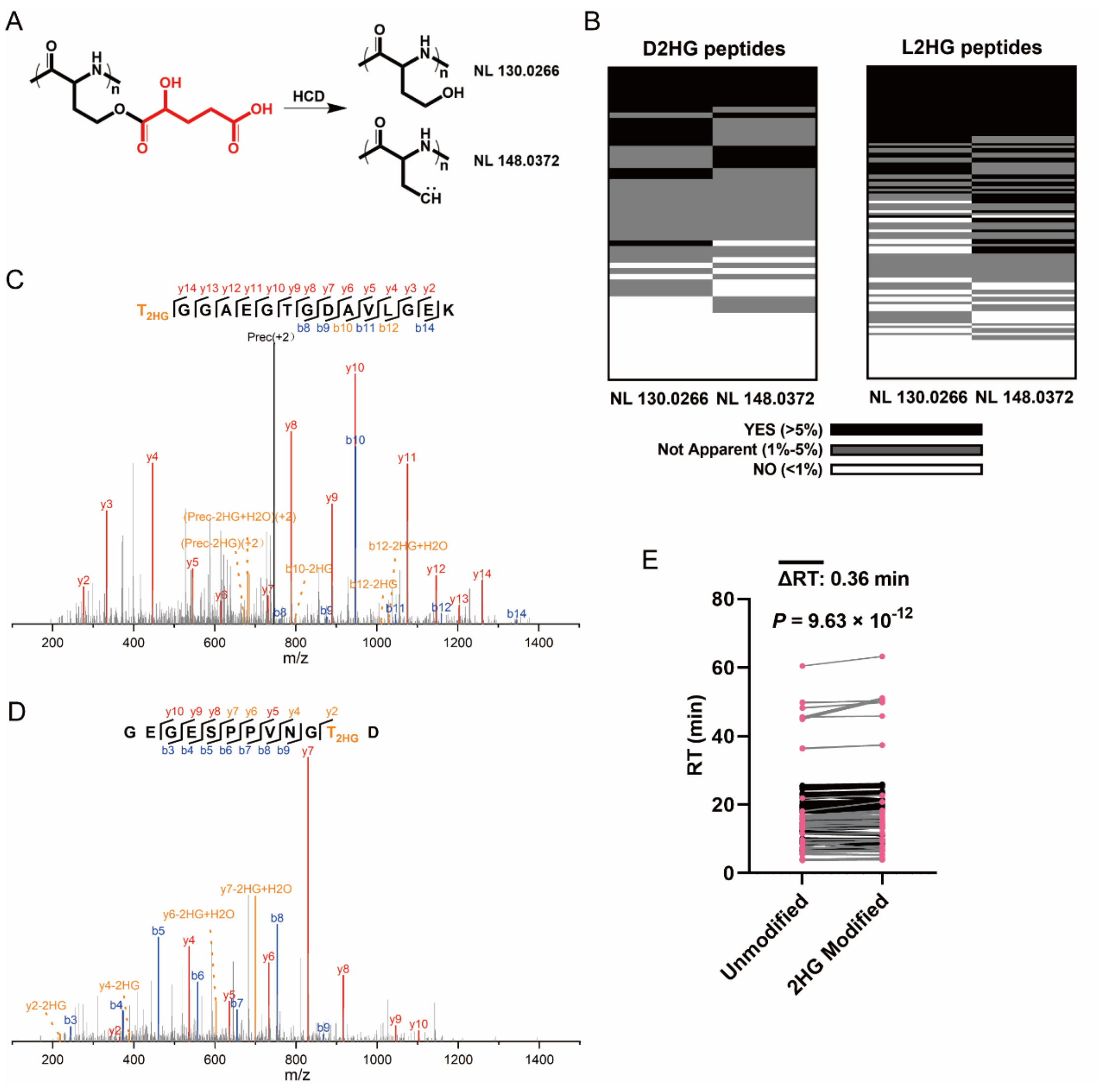


**Fig. S9. Chromatography and MS/MS fragmentation behaviour analysis.** (A) The structure of neutral loss ions generated by 2HG modified peptides under HCD. (B) Majority of the assigned 2HG modified peptides generated neutral loss ions in HCD fragmentation. (C-D) Representative MS/MS spectra of the 2HG modified peptides. The neutral loss ions are highlighted in orange, the b ions are shown in blue and the y ions are shown in red. (E) Pairwise comparison of the retention time of 2HG modified and non-modified peptides. The *P* value was calculated by a paired two-tailed Student's t-test.

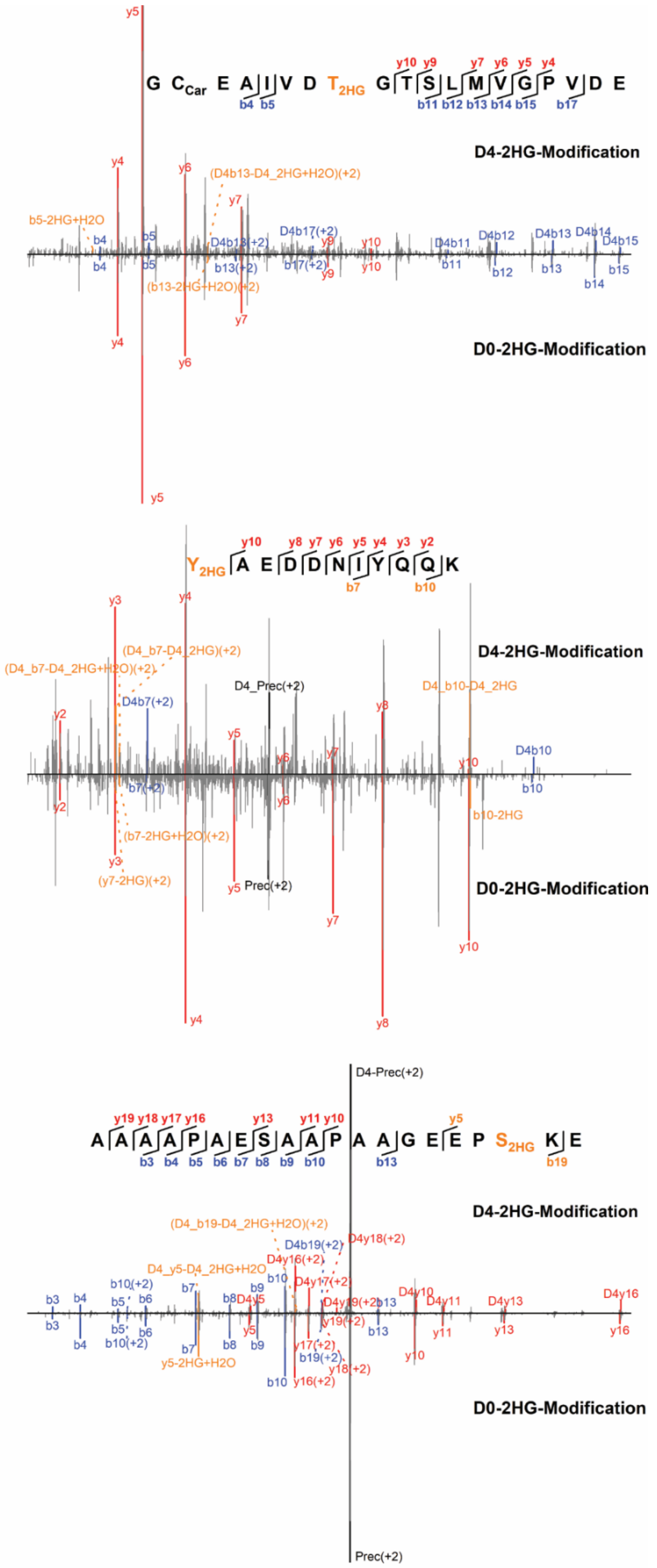


**Fig. S10.** Representative MS/MS spectra of 2HG modified peptides and d4-2HG isotopically labelled versions.

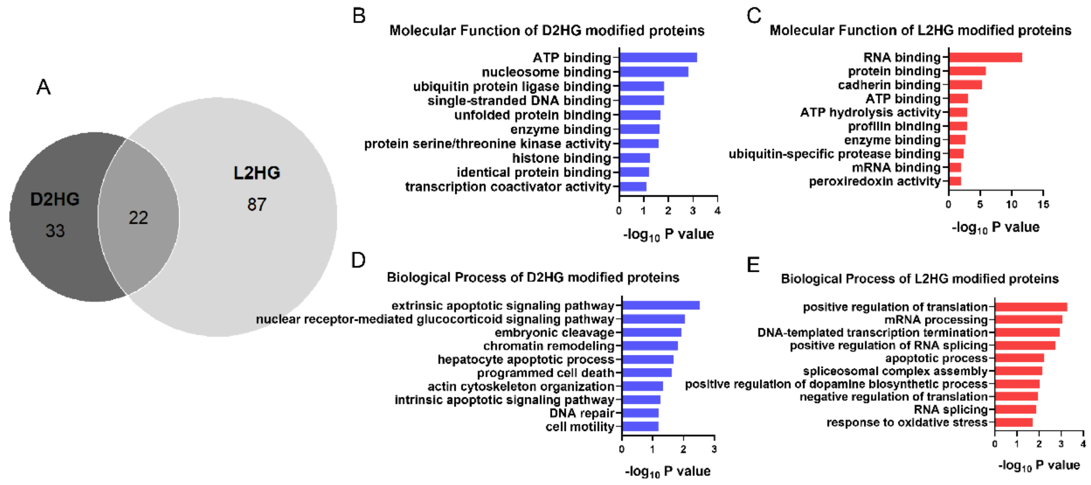


**Fig. S11. Analysis of D and L2HG modifications.** (A) Overlapped analysis of D2HG and L2HG modified peptides. (B-E) GO analysis of D2HG and L2HG modified proteins in terms of biological process and molecular function.
